# Influence of mowing timing on the breeding success of an endangered ground-nesting migratory bird, Whinchat (*Saxicola rubetra*)

**DOI:** 10.1101/2025.07.16.665079

**Authors:** Aimée Gray, Grace Walsh, Eva de la Peña, Alex S. Copland, Barry J. McMahon

## Abstract

Ground-nesting birds in Europe are declining due to anthropogenic pressures that reduce breeding productivity. Agricultural intensification under the Common Agricultural Policy in Europe is a major driver of this for farmland species, including Whinchat (*Saxicola rubetra*), in Ireland. We collected data over five breeding seasons in the Shannon Callows, a stronghold for this species where seasonal flooding limits agricultural intensification. We recorded habitat characteristics, breeding activity, and phenological events to determine population growth rates and assess mowing impacts on breeding success. Mean fledging date was July 4^th^ (± 9.28 days). We calculate that for the population to remain stable, with 80.6% brood survival, mowing should be delayed until July 14^th^ (+ 9.28 SD days to allow for natural variation). It is well documented in the literature that early mowing significantly impacts Whinchat productivity. However, when modelling mowing date against Whinchat breeding success weak significance was found that disappeared when increasing sample size through simulations, suggesting that mowing timing alone is not a strong determinant of reproductive success. Other factors including habitat structure, prey availability, predation risk, and weather conditions likely interact to influence breeding outcomes and warrant further study. We present a valuable framework with five years of breeding data that can inform Whinchat conservation and highlight the need for longer-term studies considering additional factors including how weather conditions affect mowing and prey availability.

## Introduction

European bird populations have suffered declines in recent decades, including substantial losses in breeding numbers (Keller et al. 2020; Burns et al. 2021). Agricultural species have undergone particularly steep declines, largely driven by the impacts of the Common Agricultural Policy (CAP) and subsequent intensification of agricultural systems, which has negatively affected specialised bird species (Donald et al. 2001; Newton 2004; Reif and Vermouzek 2019; Rigal et al. 2023). Agricultural land is the most widespread land use in Europe occupying approximately 40% of the land surface area (Eurostat 2021) and ground-nesting birds in these landscapes are particularly vulnerable and at risk of decline (Bas et al.2009; McMahon et al. 2020; McMahon et al. 2024). Examples include Corncrake *Crex crex* (Green and Stowe 1993), Skylark *Alauda arvensis* (Krebs et al. 1999; Donald et al. 2002), and Whinchat *Saxicola rubetra* (Müller et al. 2005; Grüebler et al. 2012; Grüebler et al. 2015a; Grüebler et al. 2015b) which have declined in population and in breeding range. The pressures on these species are typically a result of agricultural intensification (Douglas et al. 2023) which leads to habitat loss (Traba and Morales 2019), increased generalist predators (McMahon et al. 2020) and changes in mowing practices (Tome et al. 2020).

Whinchat *Saxicola rubetra* are an Afro-Palearctic migrant that have declined in population and breeding range across Europe (Müller et al. 2005; Grüebler et al. 2012; Grüebler et al. 2015a; Grüebler et al. 2015b). They are considered an indicator species of low intensity, high nature value farmland and land-use changes (Britschgi et al. 2006; Orłowski et al. 2017). Overwintering habitat conditions have minimal impact on the species’ survival (Hulme and Cresswell 2012; Blackburn and Cresswell 2015). Mortality during migration, particularly for first year birds may be impacting populations (Blackburn and Cresswell 2015), however migration barriers, present during spring migration at least have not been shown to effect behaviour of juveniles compared to adults (Blackburn et al. 2019). The issues with migration are likely due to the quality and availability of stopover sites and further study is needed to address this (Cresswell et al. 2025). Nonetheless, survival at breeding grounds remains a problem indicating that European conservation measures are urgently required. The European breeding population is at risk with a ten-year trend of −33% (2014-2023) (PECBMS 2025). In Ireland, a 76% decline in Whinchat breeding habitat has red-listed the species and designated it as high conservation priority (Balmer et al. 2013; Gilbert et al. 2021). The Irish population is estimated at 51-100 pairs (Crowe et al. 2021). The species has two strongholds remaining in Co. Wicklow in the east the Shannon Callows in the Midlands (Balmer et al. 2013).

Whinchat use botanical meadow grassland (Fischer et al. 2013), which was once widespread in many European countries, but is now disappearing with the continued transition to intensive agriculture. In Ireland, meadow habitat is restricted to a few pockets of unimproved agricultural land (Fossitt 2000). The majority of farmland in Ireland is improved grassland associated with dairy farming which is less likely to be associated with the participation in agri-environmental schemes (Sheridan et al. 2011; Sheridan et al. 2017). Whinchat require high insect production, varied vegetation and sufficient perches (Murray et al. 2016). As they nest on the ground, they can be impacted by mowing practices (Tome et al. 2020) and therefore they also require sufficient cover until birds can fly. After leaving the nest, chicks will remain hidden in the grass for ∼7 days being fed by their parents, (Collar 2005; Tome and Denac 2012; Storchová and Hořák 2018) and during this time are still vulnerable to mowing. Changing conditions, associated with changing mowing regimes, on breeding grounds (Vickery et al. 2014) are considered the biggest threat to the Whinchat. The Shannon Callows, a seasonally flooded grassland area along the banks and islands of the River Shannon, is such an unimproved agricultural grassland. Mowing in the Shannon Callows has historically been highly dependent on weather conditions (Heery 1993), and staggered throughout the area due to different land ownership (Martin et al. 2023). Mowing regimes have also been controlled to a degree since 1993 through grassland management measures of various agri-environmental schemes for Corncrake conservation (Copland et al. 2012). The Shannon Callows are designated as a Special Protection Area for the protection of breeding Corncrake *Crex crex*, and as such were included in agri-environmental schemes to control mowing for this species (Copland et al. 2012), which would have benefitted Whinchat. Corncrake are now locally extinct in the area meaning many hay meadows cannot be included in these schemes.

An important role of restructured agri-environmental policy is to halt the decline of European bird populations. For this, species specific measures implemented over a European spatial scale has been recommended (Broyer et al. 2014b). This research will focus on strengthening the scientific backing for the creation of a Whinchat agri-environmental measure (AEM), which should be implemented at the continental level before declines worsen. There are AEMs in Europe which include controlling and postponing mowing (Broyer et al. 2014a) and the provision of fallows to create perches (Küblbeck et al. 2024). The control of grazing and stocking densities is another candidate AEM which could impact Whinchat breeding success (Murray et al. 2016; Calladine and Jarrett 2021). Studies suggest hay mowing be delayed until, at least, July 1^st^ in Europe to protect meadow passerine populations against extinction (Fischer et al. 2013; Broyer et al. 2017). Studies from Ireland have suggested July 27^th^ as a suitable mowing date although this was a pilot study based on a small sample size over one year (Kenny 120 et al. 2015).

The current study aims to update previous recommendations (Kenny et al. 2015) with an additional four years of data for the Shannon Callows. This study aimed to characterise the breeding activity of Whinchat in the Shannon Callows, to determine a safe mowing date to support a sustainable population. As ground-nesting birds, Whinchat are particularly vulnerable to mowing practices. By analysing five years of breeding data, we aim to provide informed conservation recommendations to help the viability of this population.

## Methods

### Study site

The Shannon Callows is a protected area of lowland wet meadows and extensively grazed pasture in central Ireland. The area is approximately 50km long with a high diversity of semi-natural habitats. Land here is managed for silage and hay and much is unsuitable for intensive agriculture. It is designated as a Special Protection Area for the protection of wintering birds, their habitat, and breeding Corncrake *Crex crex* under the Birds Directive (2009/147/EC) and as a Special Area of Conservation under the Habitats Directive (92/43/EEC). The extent of the Special Protection Area is shown in Fig 1.

**Fig. 1.**
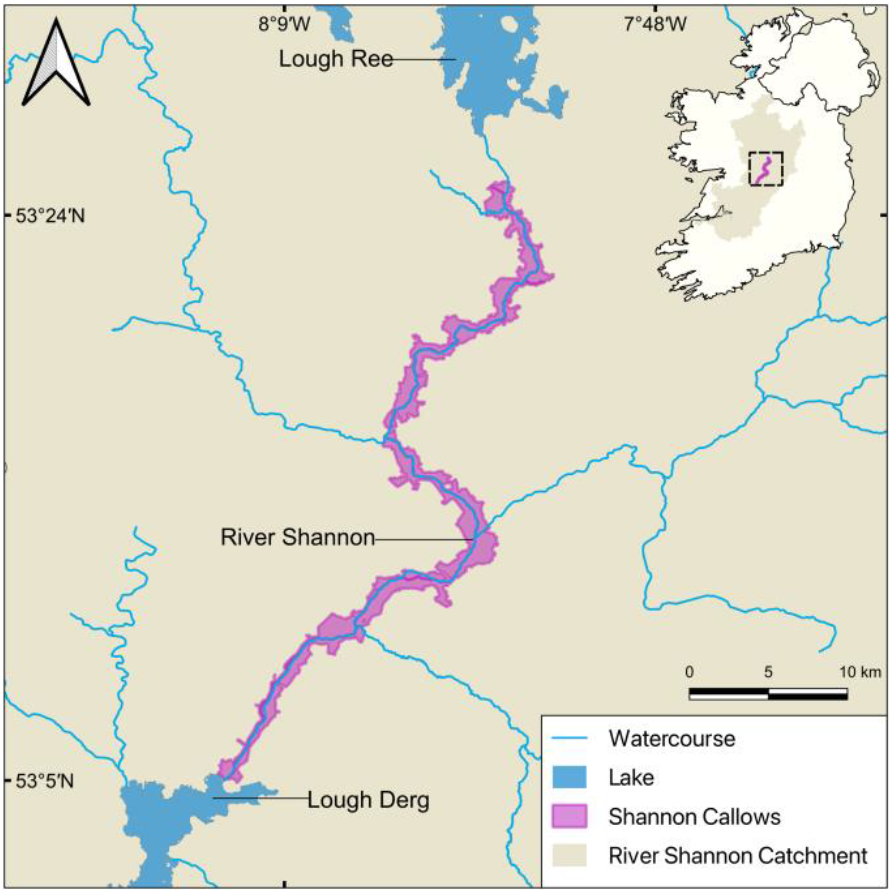
Location of the Shannon Callows in central Ireland where the study was undertaken.Data: NPWS, Special Protection Areas (SPA), 2024. CC BY 4.0.

### Study species

Whinchat breed in the Shannon Callows, which is one of their strongholds in Ireland (Balmer et al. 2013). Chicks typically leave the nest at 13.66 days (± 0.85 SD) (Storchová and Hořák 2018; BTO 2023), but in some places up to 15 days old (Tome and Denac 2012). They can fly at days 17-19, but after this they still remain near the nest and dependent on their parents for 9-15 more days with independence around day 29-30 (Collar 2005; Tome and Denac 2012; Storchová and Hořák 2018). Up until day 30, fledglings remain within approximately 100 meters of the nest site (Tome and Denac 2012) allowing them to be reliably assigned to specific nests. In this study we record fledging date once fledglings are observed. As they remain on the ground or in the nest for several days after fledging our date is likely overestimated. Our focus was on when chicks had sufficient mobility to avoid mowing and therefore fledging date in this study can be defined as such rather than the strictly biological interpretation.

### Field methods

We collected field data on breeding whinchat in the Shannon Callows in 2014, 2016, 2018, 2019 and 2020. In 2018 a comprehensive Whinchat census was carried out across all suitable habitats. Subsequently, in 2019 and 2020, surveys were limited to previously confirmed sites. For 2014 and 2016, only fledging date data were available. Surveying was conducted at a slow pace, with transects spaced 50-100m apart, depending on accessibility. In areas where drains or ditches restricted movement, potential perch points such as trees, fences and plant stalks were scanned for bird activity. Nest disturbance was minimised throughout the survey period.

Data collection focused on habitat characteristics, breeding activity and phenological events. Key recorded dates included pair confirmation, first and last observations, breeding, fledging, and mowing. The variables used in the analysis are shown in Table 1. Nests were considered successful if they produced fledglings. To standardize temporal data, date was converted to ordinal day, accounting for leap years.

**Table 1.** Definitions of terms used in the study to model and assess breeding Whinchat success in the Shannon Callows, Ireland in relation to timing of mowing.

| Term | Description |
| --- | --- |
| Breeding date | Date adult observed carrying food OR both adults observed alarm calling (both indicate hatched young) |
| Fledging date | Date young were first observed in territory (recently fledged birds will hide in grass so likely overestimated). Used to indicate when birds could avoid mowing activity. |
| Predicted hatched date | Estimated from breeding date |
| Final status | Success – bird fledged; Failure – breeding pair were recorded but no fledglings |
| Mowing day | Date mowing was observed within that nest's estimated territory |
| Predicted Fledging Date | Predicted hatching day plus 15 days (assumed maximum time chicks take to fledge) |
| Mowing Day in Relation to Fledge Timing | Difference between the predicted fledging date and the mowing date |

Surveys were conducted from April to August, with site visits rotated to ensure comprehensive coverage. Observations took place between 06:00–12:00 and 16:00–20:00 Irish Standard Time (IST). During this period, sunrise ranged from approximately 07:00 in April to 05:00 in June, and sunset from 20:00 in April to 22:00 in June. At the beginning of the breeding season, males sing from perches, allowing both visual and auditory detection. Male perch locations were recorded using GPS coordinates, with behavioural observations made from approximately 80 meters to minimize disturbance. A second observer confirmed territory boundaries each June to ensure accuracy. Territories were classified following Bibby *et al*., (2000), requiring a minimum of three observations over a 10-day period. Once pairs were established, spot mapping was used to estimate territory boundaries (Bibby 2000). Territories were monitored every 3-5 days for 10-20 minutes to record key breeding behaviours, including pairing (birds observed perching together), breeding activity (alarm calling, carrying food), and fledging success (direct observation of fledglings).

### Cumulative proportions

The population growth rate (λ) was calculated with 188

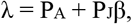

where P_A_ is the probability of annual adult survival, P_J_ is the probability of juvenile survival from fledging to the following breeding season, and β is the production of female fledglings per pair per breeding season (Pulliam 1988). When λ equals 1, population size remains constant, when it is greater than 1 the population is increasing and when it is less than one the population is decreasing (Pulliam and Danielson 1991) (see Table 2 for justification of estimates). A literature review was undertaken to identify these whinchat vital rates. 197

**Table 2.** Vital rates used to calculate the population growth rate (λ) with justification and sources for each.

|  | Value | Justification | Literature |
| --- | --- | --- | --- |
| Probability of annual adult survival ($P_A$) | 0.52 | Can be considered a true estimate of survivability due to highly site faithful birds in Nigeria | (Blackburn and Cresswell 2016) |
| Probability of juvenile survival ($P_J$) | 0.276 | Mean of 3 estimates | (Schmidt and Hantge 1954; Müller et al. 2005; Border et al. 2017) |
| Breeding productivity ($\beta$) | 2.61 | Mean of 8 estimates (5.21/2) | (Gray 1973)(5.6); (Fuller and Glue 1977)(4.96); (Pudil 2001)(5.48); (Gritschik and Baranovsky 2004)(4.64); (Müller et al. 2005)(4.9); (Frankiewicz 2008)(5.6); (Fischer et al. 2013) (4.73); (Shitikov et al. 2015)(5.79) |

### Statistical modelling and resampling

First, we created a new variable called *Predicted Fledge Date* under the assumption that chick fledging occurs 13.66 (± 0.85 SD) (Storchová and Hořák 2018; BTO 2023) days following the *Predicted Hatching Date*. This is the stricter biological definition of fledging, referring specifically to when chicks leave the nest. Given the absence of data regarding the *Predicted Hatching Date* for the 53 available observations, the dataset was subsequently reduced to 43 observations for further analysis. Subsequently, a second new variable, *Mowing Day in Relation to Fledge Timing*, was calculated.

This variable was calculated as follows:

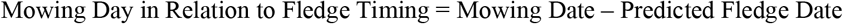

Mowing Day in Relation to Fledge Timing = Mowing Date – Predicted Fledge Date 224 Thus, for this new variable (*Mowing Day in Relation to Fledge Timing*) a negative value indicates that mowing occurred prior to the chicks fledging, while a positive value denotes that mowing took place after the chicks had left the nest.

A generalized linear model (GLM) with a binomial error distribution was applied to assess the relationship between Whinchat breeding success and timing of mowing relative to fledging. This analysis was performed using the *glm()* function from the stats package in R. The significance of the predictor variable was evaluated using Wald’s z-tests.

To further assess the robustness of the results, three complementary statistical approaches were carried out. First, bootstrapping was carried out, where the variability of model coefficients was estimated using a resampling procedure with 1000 replicates, implemented via the *boot* package (Davison and Hinkley 1997). Confidence intervals for the model estimates were computed using the percentile method. After that, a five-K-fold cross-validation procedure was conducted using the *caret* package (Kuhn 2020) to evaluate model predictive performance. The root means square error (RMSE), means absolute error (MAE), and R^2^ were used to assess the model’s explanatory power. Finally, we also performed Monte Carlo Simulations to explore model sensitivity to sample size. Thus, 10000 simulated observations were generated based on the distribution of existing data. The impact of increasing the sample size on coefficient estimates and statistical significance was examined using the *simglm* package (LeBeau 2022).

Plots were generated using the *ggplot2* package (Wickham 2016) in R. All statistical analyses were conducted using RStudio (Team 2024) and were performed at a significance level of α =248 0.05.

## Results

### General description of data

Over five survey years in the Shannon Callows a total of 181 Whinchat territories were monitored, consisting of 157 breeding pairs and 24 single males. Between 2018 and 2020, 68% of nests successfully produced fledglings. In 2018, 19 pairs (68%) fledged young, followed by 35 (69%) in 2019 and 29 pairs (66%) in 2020. The earliest recorded fledging date was on the 9^th^ of June 2016 while the latest was the 28^th^ of July 2019. On average, fledging occurred on July 2^nd^. Since fledging date was recorded based on the first observed presence of young birds, these dates are likely over estimated. A summary of the total number of nests per year and the ordinal day of fledging is provided in Table 3.

**Table 3.**
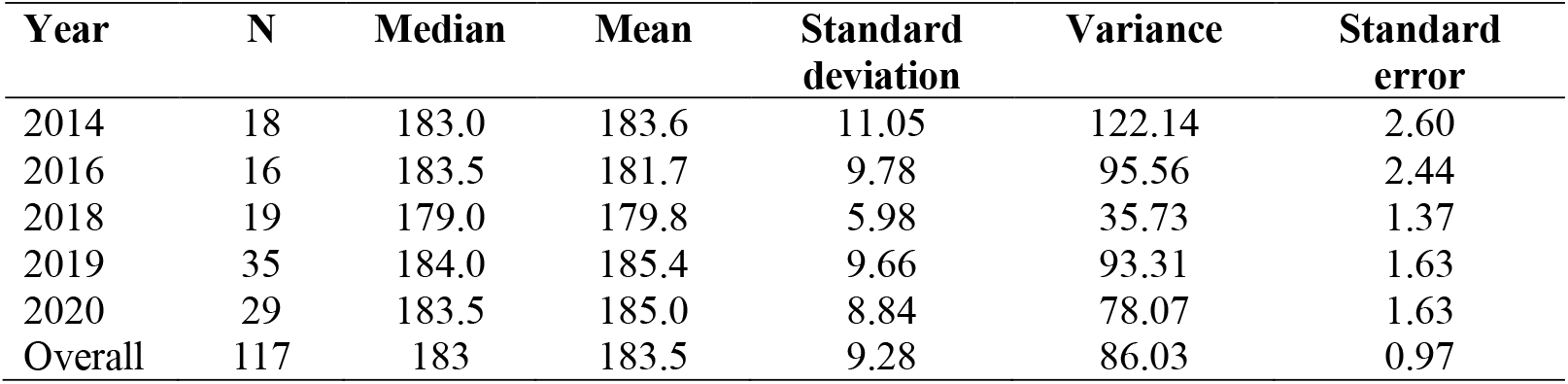
Summary statistics of Whinchat fledging dates in ordinal day (OD) for the study years, including sample size (N), median, mean, standard deviation, variance, and standard error of the mean.

| Year | N | Median | Mean | Standard deviation | Variance | Standard error |
| --- | --- | --- | --- | --- | --- | --- |
| 2014 | 18 | 183.0 | 183.6 | 11.05 | 122.14 | 2.60 |
| 2016 | 16 | 183.5 | 181.7 | 9.78 | 95.56 | 2.44 |
| 2018 | 19 | 179.0 | 179.8 | 5.98 | 35.73 | 1.37 |
| 2019 | 35 | 184.0 | 185.4 | 9.66 | 93.31 | 1.63 |
| 2020 | 29 | 183.5 | 185.0 | 8.84 | 78.07 | 1.63 |
| Overall | 117 | 183 | 183.5 | 9.28 | 86.03 | 0.97 |

Mowing data was also collected from 2018 to 2020. Across this period, 54 sites were mowed with 20 failed nests and 33 successfully fledging. In contrast, 69 sites remained unmowed, with 20 nests failing and 49 successful fledging.

### Cumulative proportions

The population growth rate (λ) was calculated as the sum of adult survival probability and the product of juvenile survival probability and the number of female fledglings produced per pair per breeding season (Pulliam 1988), equalling 1.240. This theoretically allows the population0020to remain unchanged even with a loss of 19.4% of broods in a breeding season (1-(1/1.24036)= 19.4%), meaning 80.6% of broods must survive. In this study, fledging was defined as the point at which chicks are first observed outside the nest (see Table 1). This is likely an overestimation of actual fledging date as chicks will remain hidden in the grass for some time. However, it is an indication of when chicks have sufficient mobility to avoid mowing which is the focus here. We plotted data (Fig 2) from five breeding seasons (n = 117) and fitted a linear regression model (y = mx + b) to determine the x-axis value in OD, corresponding to 80.6%. This was

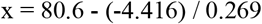

with x = 194.4, or the 14^th^ of July. This has been rounded up to the 14^th^ of July (ordinal day 195) to identify the earliest full day the threshold had been reached.

**Fig. 2.**
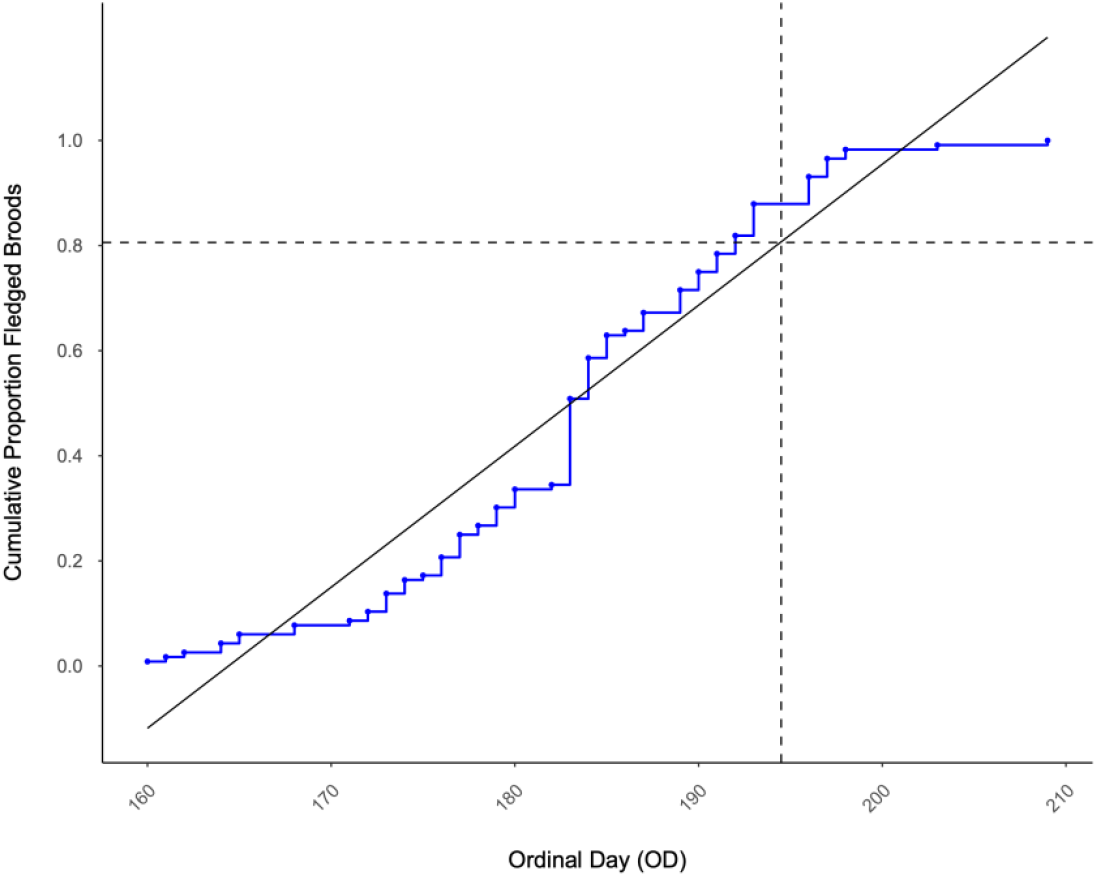
Cumulative proportion of mobile Whinchat broods by date as ordinal day (OD) for 5 years of Whinchat breeding data in the Shannon Callows, Ireland with a regression line fitted.The intercept was calculated to identify the date where 80.6% of nests were mobile (indicated by the dashed line). This is the 14th of July.

### Mowing timing effect in Whinchat breeding success

The GLM results showed that mowing timing relative to fledging was a significant predictor of breeding success (β = 0.099, SE = 0.051, z = 1.976, *p*-value = **0.048**), indicating a positive association between mowing occurring later in the season and fledging success (Fig 3).

**Fig. 3:**
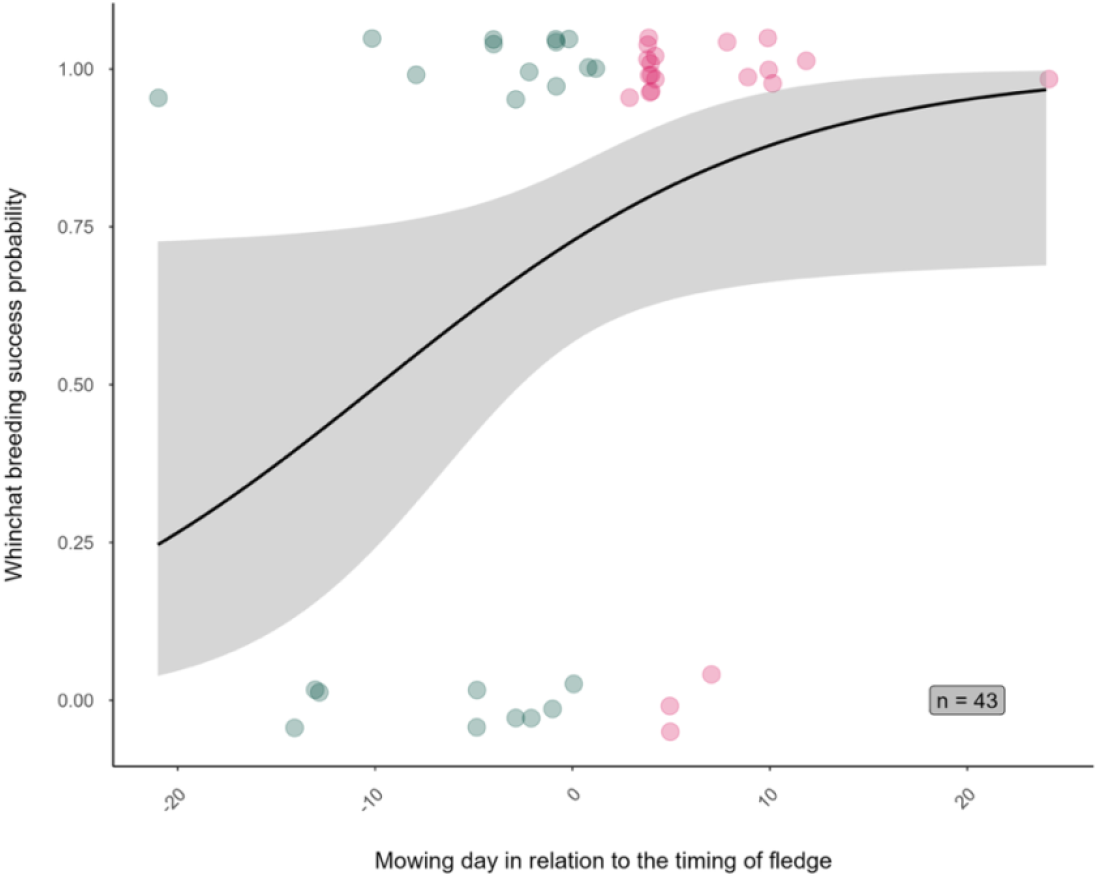
Breeding success probability of Whinchat in relation to mowing date relative to fledging timing in The Shannon Callows (Ireland, n= 43 nests). The x-axis represents the number of days between mowing and fledging (negative values indicate mowing occurred before fledging, positive values indicate mowing occurred after fledging). The y-axis shows the predicted probability of breeding success derived from a GLM (solid black line), with a 95% confidence interval (shaded area) estimated using a smoothing function. Raw data points are shown, with green dots indicating mowing events before fledging and pink dots indicating mowing events after fledging.

The bootstrapping analysis supported the stability of the model estimates. The bootstrap bias was small (0.016 for the mowing timing coefficient), and the 95% confidence interval (0.015, 0.270) did not include zero, indicating a statistically significant relationship.

The cross-validation results indicated moderate predictive performance, with an RMSE of 0.454, MAE of 0.386, and R^2^ = 0.218. This suggests that mowing timing alone explains only 21.8% of the variability in breeding success, highlighting the need for additional explanatory variables to provide greater clarity on the breeding success of Whinchat based on our dataset. However, Monte Carlo simulations revealed that when sample size was artificially increased to 10000 observations, the significance of the mowing timing variable disappeared (β = 0.004, S.E. = 0.003, *z* = 1.448, *p*-value = 0.148). This result suggested that with a larger dataset, the observed effect may not be biologically meaningful and that other ecological factors are likely to influence breeding success.

## Discussion

Data from other studies demonstrate that mowing impacts on Whinchat breeding success are significant (Tome et al. 2020). However, the full magnitude of its effect in an Irish context has not been fully investigated except with data from a single year (Kenny et al. 2015) and given this population is at the western extent of the breeding range, representing an ecological niche, we feel these new data are important. In this study we use five years of data to study the characteristics of Whinchat breeding in the Shannon Callow in Ireland. Using vital rates from the literature along with data we collected on fledging date, we calculated a mow date that would allow the population to remain stable. We also directly test for the effect of mowing in the Shannon Callows and find a statistically significant relationship between mowing timing and breeding success. However, the significance was lost when sample size was increased through simulations. Therefore, it appears other variables are impacting Whinchat breeding success in the Shannon Callows and this study therefore provides important insights for conservation applications. Additional research into these other variables for example weather, predation, food availability and vegetation would be beneficial. Nonetheless, it is still likely that if early mowing occurs (Tome et al. 2020) in a large enough area this would significantly impact on Whinchat breeding success in the Shannon Callows.

Using previously published Whinchat vital rates from across their breeding range for the probability of juvenile survival (UK (Border et al. 2017), Switzerland (Müller et al. 2005) and Germany (Schmidt and Hantge 1954)), breeding productivity (UK (Gray 1973), (Fuller and Glue 1977), Czech Republic (Pudil 2001), Belarus (Gritschik and Baranovsky 2004), Switzerland (Müller et al. 2005), Poland (Frankiewicz 2008), Germany (Fischer et al. 2013) and European Russia (Shitikov et al. 2015)) and one wintering population in west Africa (Blackburn and Cresswell 2016), we estimated the percentage loss of Whinchat broods that could, theoretical, maintain a stable population. Currently, such demographic data are unavailable for the Irish Whinchat population. Therefore, our objective is to provide a conservation framework rather than a definitive implementation strategy. Should estimates of vital rates for the Irish population become available in the future, they can be readily incorporated into this model. It is also important to acknowledge that apparent survival may differ significantly from true survival in Whinchat populations (Shitikov et al. 2015), and potentially between sexes (Müller et al. 2005; Grüebler et al. 2008), meaning that our calculations should be considered informed estimates rather than precise predictions. We calculated the population growth rate as 1.24, indicating that the population could theoretically remain stable with a loss of 19.4 % of broods in a breeding season which is equivalent to a survival rate of 80.6%. By fitting a linear regression to five years of fledging data, we estimated that 80.6% of chicks had fledged by July 14^th^ (OD = 204). This date represents when mowing could theoretically begin without major impact on Whinchat breeding success. We defined fledging as the first observation of chicks out of the nest, which may slightly overestimate true fledging but reflects when they can likely avoid mowing. To account for natural variation, we added one standard deviation (9.3 days) to the mean threshold date, yielding July 23rd. Therefore, mowing should ideally be delayed until after July 23^rd^ to reduce risk to late fledging chicks.

There are three feasible ways for this to be interpreted. Firstly, choose 80.6% of nests and protect with delayed mowing until all chicks have fledged. This does not account for stochastic processes which may reduce survival such as predation, adverse weather or sample outliers. Another option would be to protect 100% of nests, until 80.6% have fledged. This is the most time consuming and expensive option and mowing will need to be delayed on the largest area. It can be difficult to be certain of nest numbers and therefore this would make it difficult to protect a certain number without high survey effort. Lastly, and what we recommend, is to protect 89.8% of broods. The options suggested, require careful consideration and further investigation by agro-economists, policy experts and conservationists. The benefits must outweigh or at least equate the costs for conservation to be sustainable. The location of protected nests is also a consideration, as nests in certain habitat types such as edge habitats have been shown to have lower productivity in other ground nesting species (Sheridan et al. 376 2020).

Although our analysis indicates a statistically significant relationship between mowing timing and Whinchat breeding success (p = 0.048), the strength of this association is relatively weak. When the sample size was increased through Monte Carlo simulations, the significance of the predictor was lost, suggesting that the effect may not be robust. While preliminary results suggest that delayed mowing may improve fledging success, the overall explanatory power of the model remains limited. This implies that mowing timing alone is unlikely to be a strong determinant of reproductive success.

Nonetheless, mowing is a widely recognized threat to Whinchat populations and deserves further investigation, particularly in our study area, the Shannon Callows. The plausible drivers of declines are breeding failures due to bird casualties (adult and/or juvenile) (Müller et al. 2005; Grüebler et al. 2008; Grüebler et al. 2012) and reduced food availability (Vickery et al. 2001; Britschgi et al. 2006). Both of which can occur from intensified grassland management and early mowing (Feehan 2003). Previous research has demonstrated that early or intensive mowing can negatively impact Whinchat survival (Tome et al. 2020), and more broadly, reduce species richness in grassland ecosystems (Canonne et al. 2024). As agricultural intensification continues to promote earlier mowing regimes, targeted conservation measures are increasingly necessary.

Furthermore, recent studies highlight that factors such as habitat structure, predation risk, and weather conditions also play significant roles in Whinchat breeding success. For instance, vegetation heterogeneity and the presence of perching sites are crucial for nesting and foraging efficiency (Murray et al. 2016). Predation risk, particularly from mammals during the early fledgling stage, can significantly affect juvenile survival (Naef‐Daenzer and Grüebler 2016). Additionally, adverse weather conditions, such as increased rainfall and temperature extremes, have been linked to reduced fledgling survival and recruitment rates (Halliwell et al. 2023).

Edge avoidance also affects ground nesting birds in Ireland (Sheridan et al. 2020) and may influence nest selection in Whinchat. Other significant pressures on the population is increasing the likelihood of Whinchat replacement brood production at specific rare locations (Müller et al. 2005), the dissolution of pair bonds with the onset of mowing (Grüebler et al. 2015b), predation pressure from winged and terrestrial predators (Tome and Denac 2012) and detrimental weather conditions with rain, in particular, affecting nestling success (Öberg et al. 2015). Given these complexities, future studies should incorporate these additional variables to better explain variation in breeding outcomes and support more effective management of Whinchat populations.

Also mowing on the Shannon Callows has historically been dependent on weather conditions (Heery 1993). Weather conditions can be particularly impactful on the Shannon Callows. Flood extent can be a determinant of the timing of breeding, particularly if there is summer flooding which can directly cause nest failure. In 2019 and 2020, there was a wet summer which reduced early mowing, and mowing was recorded to have directly affect much less nests. In comparison in 2018 there was a very wet spring followed by a dry summer which are normal conditions. Had the summer of 2019 and 2020 been more typical, there may have been more mowing and a more significant impact on Whinchat.

Controlling mowing operations is still required and may improve Whinchat fledging success (Grüebler et al. 2012; Broyer et al. 2014b; Denac 2015; Horch and Spaar 2015; Siems-Wedhorn 2015). Meadow passerine populations in Europe have responded well with even short-lived delayed mowing projects across Europe (Broyer et al. 2016). Ireland is in a unique position to pursue the restoration of Whinchat populations on The Shannon Callows, a largely undisturbed 50km linear stretch of pasture and wet meadow landscape. Delayed machinery practice has previously been successfully implemented to improve the numbers of corncrakes in Ireland (McDevitt and Casey 2004), demonstrating merit for future conservation measures on the Shannon Callows. 434 While our results indicate that delayed mowing may enhance Whinchat fledging success, the overall explanatory power of the model remains limited, and the statistical significance of mowing timing diminishes under larger simulated sample sizes. This suggests that mowing timing alone is unlikely to be a primary driver of reproductive success in this species. A more comprehensive understanding of breeding outcomes in our study area will likely require the integration of additional ecological variables. Future research should consider incorporating factors such as habitat structure, prey availability and hunting opportunity, predation risk, and weather conditions, all of which are known to interact with land management practices and influence reproductive performance in this species.

## Acknowledgments

This research has been supported by the Irish Research Council, (Grant/Award Number: ‘GOIPG/2017/1028’). The authors gratefully acknowledge the BirdWatch Ireland Banagher team, Seán Kelly and Shane Cully, along with Kieran Kenny for their expert advice and fieldwork assistance. 451

## Statements and Declarations

The authors have no conflicts of interest to declare.

Author contributions: Aimée Gray contributed to the study conception and design, sourced funding, carried out data collection and commented on previous versions of the manuscript.

Grace Walsh wrote the manuscript and inputted into the statistical analysis.

Eva de la Peña carried out the statistical analysis and commented on previous versions of the manuscript.

Alex S. Copland contributed to the study conception and design, sourced funding, helped with data collection and commented on previous versions of the manuscript.

Barry J. McMahon contributed to the study conception and design, sourced funding, inputted into the statistical analysis and commented on previous versions of the manuscript.

All authors read and approved the final manuscript.

## Data availability statements

Data is available on request

